# Bayesian inference reveals a complex evolutionary history of belemnites

**DOI:** 10.1101/2022.08.22.504746

**Authors:** Kevin Stevens, Alexander Pohle, René Hoffmann, Adrian Immenhauser

**Affiliations:** Institute for Geology, Mineralogy, and Geophysics, Ruhr University Bochum, Germany; Palaeontological Institute and Museum, University of Zurich, Switzerland

**Keywords:** belemnites, phylogeny, Bayesian inference, Mesozoic, coleoid cephalopods

## Abstract

Belemnites are an extinct group of Mesozoic coleoid cephalopods, common in Jurassic and Cretaceous marine sedimentary rocks. Despite their significance, their total group phylogeny has rarely been considered in recent decades. In contrast, most researchers restricted the assignment of families to one of the two usually recognized subgroups, the Belemnitina and the Belemnopseina. As for many fossil cephalopods, researchers have been reluctant to employ modern phylogenetic methods to illuminate belemnites’ evolutionary history.

To overcome the “dead end” of belemnite systematics, we performed the first tip-dated Bayesian analysis of belemnite phylogeny. In our analysis, the Aulacoceratida are found as the monophyletic sister group to belemnites. The Sinobelemnitidae are resolved as paraphyletic and fall outside the Belemnitina and Belemnopseina, which make up the remaining belemnites. Belemnitina is restricted to Jurassic species with generally no or apical furrows. Holcobelidae are the earliest branching Belemnopseina. Cylindroteuthids *sensu lato* (including Oxyteuthidae) are nested within Belemnopseina, contrary to the common hypothesis placing them within the Belemnitina. Duvaliidae and Dicoelitidae are recovered as members of the Belemnopseina, but their precise relationship has to be evaluated based on more taxa and additional characters. We introduce the well-supported unranked clade Pseudoalveolata, which includes Dimitobelidae, Belemnitellidae, and members of the paraphyletic “Belemnopseidae”.

The phylogeny presented here, based on reproducible and quantitative methods, contrasts with the usually applied authoritative “stratophenetic” approach to belemnite systematics, based on the overemphasis of single characters. This result is considered the basis for future studies on belemnite phylogeny, allowing for a rigorous testing of evolutionary hypotheses.

**PLAIN LANGUAGE SUMMARY:** Belemnites were common extinct cephalopods that were closely related to today’s squid and cuttlefish. The most common fossil remains of belemnites are bullet-shaped calcitic “cones” (rostrum) that cover their internal shells. Belemnites’ evolutionary history is not well known. Our study revealed an evolutionary tree of belemnites based on the statistical analysis of morphological features of the rostrum and calibrated to the known geological ages of the studied belemnite species. This approach was for the first time applied to belemnites and changed several aspects that were believed about their evolution.

## INTRODUCTION

Belemnites (Belemnitida) are an extinct group of stem-decabrachian coleoids (e.g., Fuchs et al., 2013; Hoffmann and Stevens, 2020). They are characterized by a calcitic rostrum, which is by far the most commonly preserved part of their internal shell. In this paper, the terms belemnites and Belemnitida are used only for these calcite-rostrum-bearing species. This definition excludes groups like the Belemnoteuthida and Diplobelida, sometimes referred to as belemnites. The paraphyletic assemblage of Belemnitida, Aulacoceratida, Belemnoteuthida, and Diplobelida is referred to as “belemnoids” in lieu of a proper understanding of their interrelationships at present (Hoffmann and Stevens, 2020). Diplobelida probably represent close relatives of crown-Decabrachia (Fuchs, 2019; Fuchs et al., 2013). Fuchs et al. (2013) regarded belemnites as sister to a group consisting of crown Decabrachia, the stem-decabrachian *Longibelus*, and the Diplobelida. In a cladistic analysis, Sutton et al. (2015) found the belemnite *Hibolithes* closely related to the coleoid genera *Phragmoteuthis* and *Belemnoteuthis*, nested within the Decabrachia crown group.

The relationship of the rostrum-bearing Aulacoceratida with other “belemnoid” groups, is at present also unclear (Keupp and Fuchs, 2014). Aulacoceratids have aragonitic rostra (also called “telum”; see Jeletzky, 1966) and differ in other morphological aspects from belemnites (e.g., Jeletzky, 1966; Mariotti et al., 2021), but their aragonitic rostrum likely represents the ancestral rostrum structure of coleoids.

The internal phylogenetic relationships of belemnites are even less clear than their relationship to other coleoids. Early subdivisions of belemnites relied on general external characteristics, mostly on the number and position of furrows, e.g., the classification of Werner (1913). Abel (1916) subdivided all belemnites into the families “Clavirostridae” and “Conirostridae” due to their early ontogenetic development. The definitions used nowadays for belemnite families go back to Stolley’s (1919) and Naef’s (1922) classifications. Jeletzky’s (1966) proposal of subdividing the calcite-rostrum bearing belemnites into apically furrowed Belemnitina and alveolar furrowed Belemnopseina has been largely followed by subsequent workers and has been virtually the only applied subdivision of belemnites higher than the family level since.

Recognition of the alveolar furrowed Triassic-Early Jurassic Sinobelemnitidae as true belemnites (Zhu and Bian, 1984; Iba et al., 2012; Niko and Ehiro, 2022) has significantly altered views on belemnite phylogeny. Apart from the Sinobelemnitidae, belemnites are exclusively known from the earliest Jurassic onwards. Earlier hypotheses of belemnite phylogeny focused on the well-known European fossil record of the group that suggested their origin during the Hettangian in the diminutive and relatively character-poor genera *Schwegleria* and *Nannobelus*, which both lack alveolar furrows. By the Early Jurassic, belemnites had reached a cosmopolitan distribution and relatively high diversity and abundance (e.g., Iba et al., 2014a, 2014b; Weis et al., 2015a). Although affected by second-order extinction events (e.g., Dera et al., 2016; Neige et al., 2021; De Baets et al., 2021), belemnites continued to be diverse during the Jurassic and early Early Cretaceous (e.g., Schlegelmilch, 1998; Mutterlose, 1988, 1998; Iba et al., 2011), with the two last occurring, disjunctively distributed families, the Boreal Belemnitellidae and the Austral Dimitobelidae, finally becoming extinct at the K/Pg-boundary (e.g., Doyle, 1992; Christensen, 1997; Iba et al., 2011).

The evolutionary history of belemnites as a whole has rarely been studied since Jeletzky (1966). While several authors speculated about the interrelationships of belemnite families (e.g., Christensen, 1997; Iba et al., 2012; Weis et al., 2012), there has been no study of their phylogenetic relationships based on modern phylogenetic methods. This pattern reflects a general tendency of researchers studying fossil cephalopods in the past (Neige et al., 2007; Bardin et al., 2014; Pohle et al., 2022).

This paper presents the first quantitative approach towards belemnite phylogeny based on Bayesian inference. The dichotomous subdivision of all belemnites into Belemnitina and Belemnopseina, as these groups are usually defined, is not supported by these results. Our findings challenge usual assumptions about the evolution of belemnites and identify parts of the belemnite phylogenetic tree that still lack resolution.

## METHODS

We selected 24 belemnite species, representative of the stratigraphic range, geographic distribution, and diversity of the whole group (Table 1) and scored them for 29 rostrum characters (Fig. 1; Supplementary Files 1, 2). Three aulacoceratid genera (including one putative genus) were also included. Although other fossil “belemnoid” coleoid groups are likely more closely related to the Belemnitida than aulacoceratids (e.g., Diplobelida, Belemnoteuthida), these do not have proper rostra (*sensu* Fuchs, 2012) and so do not contribute to the resolution of internal relationships of belemnites, whose phylogeny is here inferred based on rostrum characters only. For the vast majority of belemnites, the rostrum is the only known part (e.g., Hoffmann and Stevens, 2020) mimicking the situation for conodonts, where inferences of their phylogenetic relationships must also be based on conodont element data only (e.g., Donoghue, 2001; Bai et al., 2022). Morphological data comes from several published sources and our own observations (Table 1). The terminology of belemnite morphology follows Hoffmann and Stevens (2020) and Stevens et al. (2022). The character matrix was compiled with Mesquite version 3.7 (Maddison and Maddison, 2021). Coding practice follows suggestions by Brazeau (2011) for morphological character coding.

**TABLE 1.**
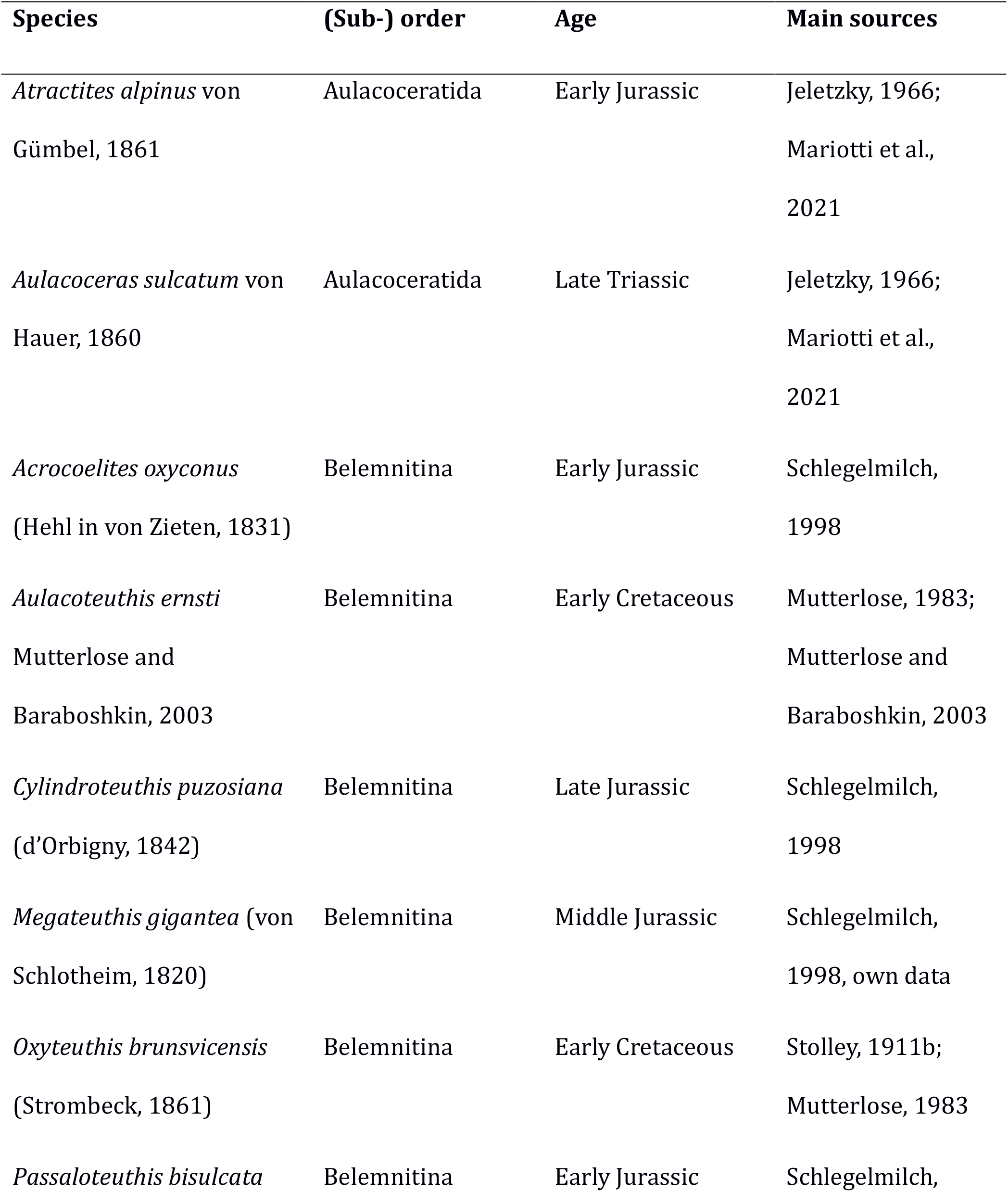
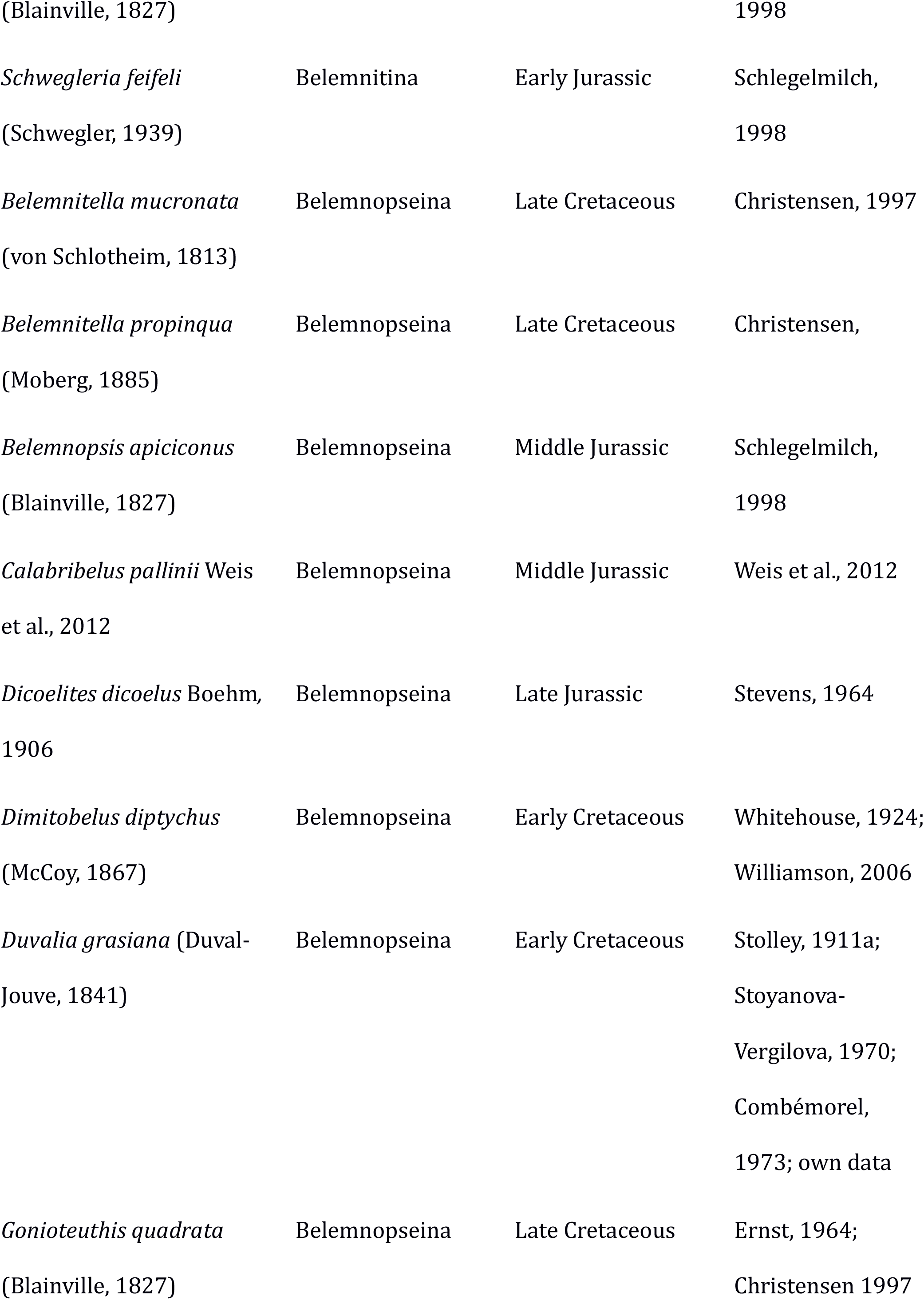
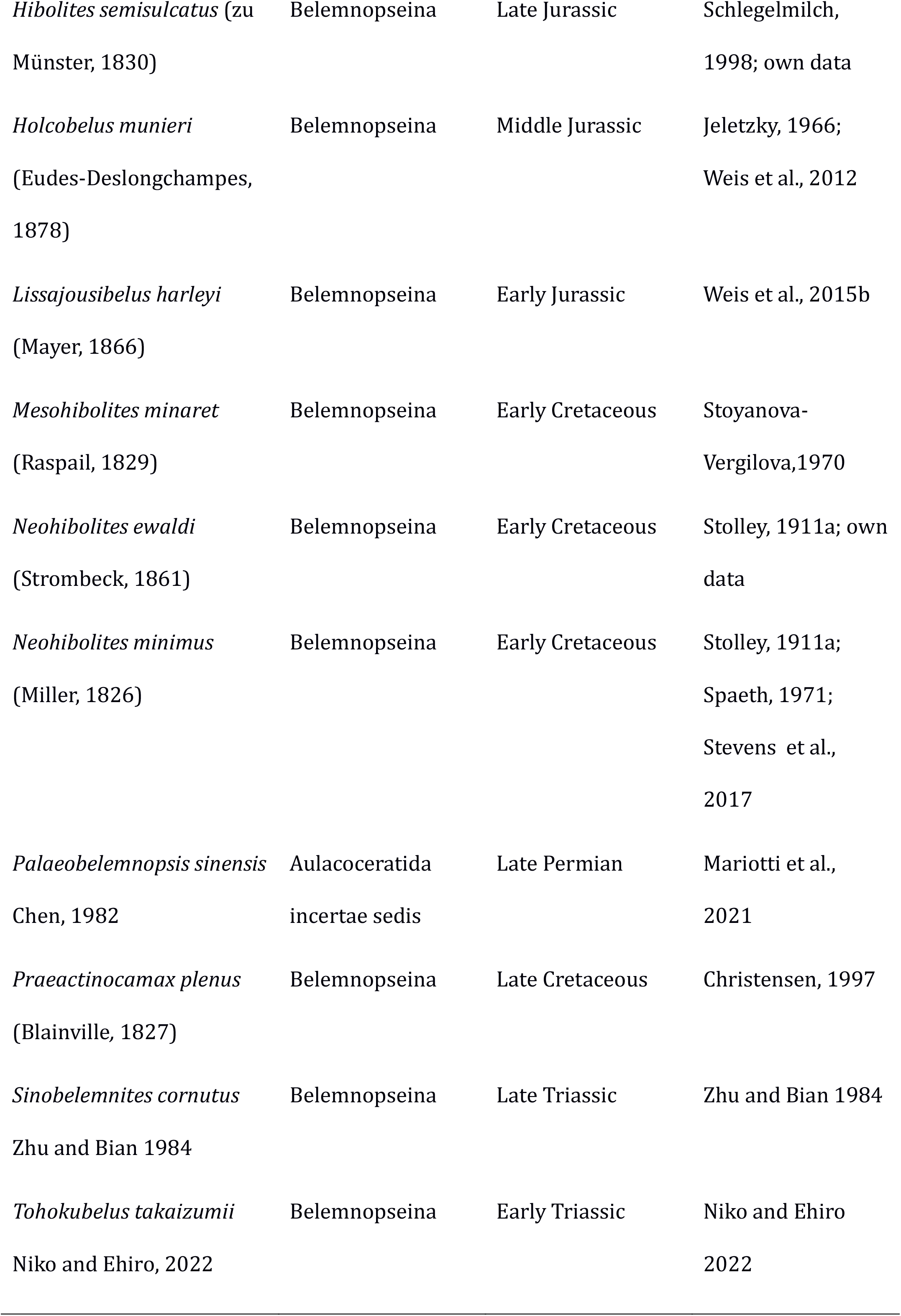
Overview of species used in the analysis. Assignment to suborders is here based on previous taxonomic concepts, which may differ from the results obtained here. For more details, see supplementary material.

**FIGURE 1.**
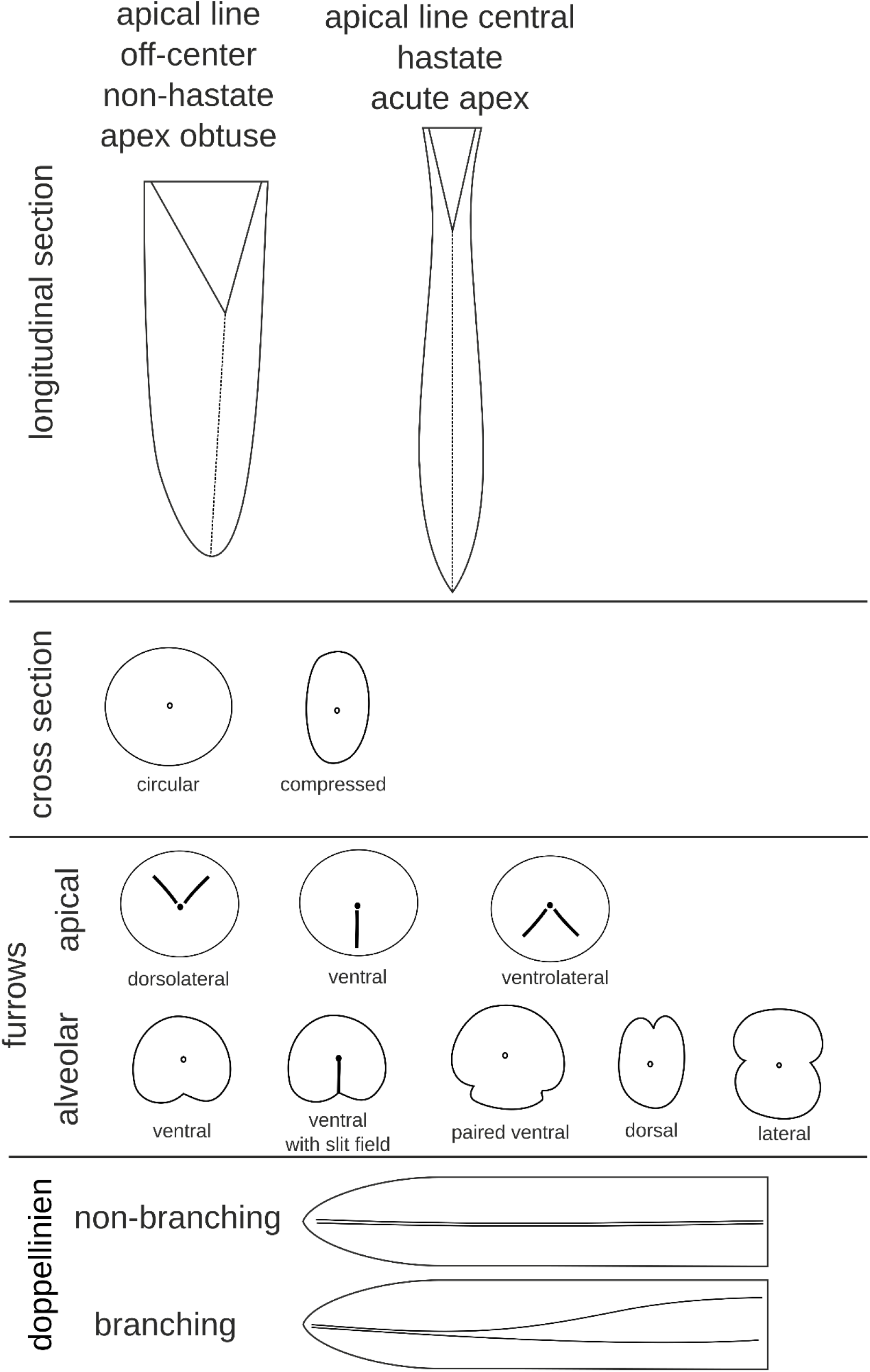
Terminology of some of the characters applied coded for the analysis (modified after Mutterlose, 1983; Doyle, 1990; Schlegelmilch, 1998; Hoffmann and Stevens, 2020).

We used Bayesian tip-dating, which has become increasingly popular in recent years for phylogenetic inference from morphological data for diverse extinct groups of invertebrates, including cephalopods (e.g., Wright, 2017; Paterson, 2019; Pohle et al., 2022). The analyses were performed in BEAST 2.6.7 (Bouckaert et al., 2019) using the fossilized birth-death model as a tree prior (Stadler, 2010; Gavryushkina et al., 2014; Heath et al., 2014) and the parametrization of net diversification rate, turnover and sampling proportion (Heath et al., 2014). Morphological character evolution was modeled with the Mkv model, including invariant site correction (Lewis, 2001). Characters were partitioned according to their number of states, except for character 13 (“Doppellinien” type); all characters were binary. The exchangeability rates were set to 1.0 for binary characters and 1.5 for the single three-state character to prevent the artificial upweighting of multistate characters (King et al., 2017), although this is naturally expected to have a minimal impact on the analyses. We furthermore accounted for heterogeneous rates across sites with two discretized gamma shape rate categories. Although usually four or more rate categories are employed in morphological datasets for this purpose (Harrison and Larsson, 2015), we used only two categories due to the small number of characters and states. Tip dates were fixed to the midpoint between the first and last occurrence date of the corresponding species. First and last occurrence dates are based on the literature and are calibrated to the ICS 2020 age model (Supplementary File 3; Gradstein et al., 2020). We used a strict morphological clock with a lognormally distributed prior (mean = 0.1, standard deviation = 1.25). We placed an exponential prior on the origin (mean = 10 my, offset = 253.1 my), limiting the youngest possible origin date to the age of the oldest taxon of the analysis. This approach avoids unrealistically old estimates while not imposing an overly informative prior. The prior on diversification rate was set to an exponential distribution (mean = 1.0), and the turnover prior to a uniform distribution between 0.0 and 1.0. For the sampling proportion, we used a uniform prior with an upper limit of 0.15, which we justify by a very rough estimate of the number of belemnite species in the Palaeobiology Database (PBDB), which resulted in c. 200 species.

Although this number likely underestimates the true number of belemnite species by some margin due to the incompleteness of both the fossil record and the PBDB, it represents a useful estimate to provide an absolute upper limit for sampling rate, as it assigns zero probability to any values above 0.15 (corresponding to the ratio between taxa used in the analysis and approximate total number of known species). Lastly, we enforced a monophyletic constraint on the Belemnitida (without the Sinobelemnitidae). We justify this constraint by the strong prior expectation that this group is monophyletic (however, note that this does not preclude potential paraphyletic relationships with respect to other groups such as Diplobelida, Phragmoteuthida, or crown-Decabrachia). The analysis was run for two separate runs of the MCMC algorithm, each with 10 000 000 generations sampling every 10 000 generations and 10% of the samples discarded as burn-in. Convergence was checked using Tracer (Rambaut et al., 2018). The tree files were combined in LogCombiner (Bouckaert et al., 2019) and the Maximum Clade Credibility tree was generated with TreeAnnotator, but using the older BEAST version 1.10.4 (Suchard et al., 2018) because, in contrast to TreeAnnotator in BEAST 2.6.7, it treats sampled ancestors as belonging to the same clade, which results in underestimated posterior probabilities (Barido-Sottani et al., 2020). The xml script to run the analysis in BEAST, the resulting combined log and tree files and the annotated summary tree are contained in Supplementary File 4.

## RESULTS

The parameter estimates of the tip-dated analyses are listed in Table 2. Although these parameters were not a focus of our current study, and more extensive model testing should be carried out before drawing any definite conclusions, they provide some insights. According to the estimated sampling rate (95% HPD interval between 0.015 and 0.15 with 27 included species), we would expect a total number of belemnite and aulacoceratid species that existed until the end of the Cretaceous to be somewhere between approximately 180 and 1 800. Although this is a rather large credible interval, it appears to be a reasonable estimate that could be refined by adding more species or occurrence data. The omission of the diplobelids and phragmoteuthids, which probably belong to the same clade, may also have caused slightly biased estimates. Furthermore, refined models with variable rates through time may also improve these estimates. The age of the last common ancestor (origin parameter) of belemnites and the aulacoceratids included here was estimated to lie within the Permian, which roughly agrees with previous hypotheses (e.g., Jeletzky, 1966; Kro ger et al., 2011).

**TABLE 2.**
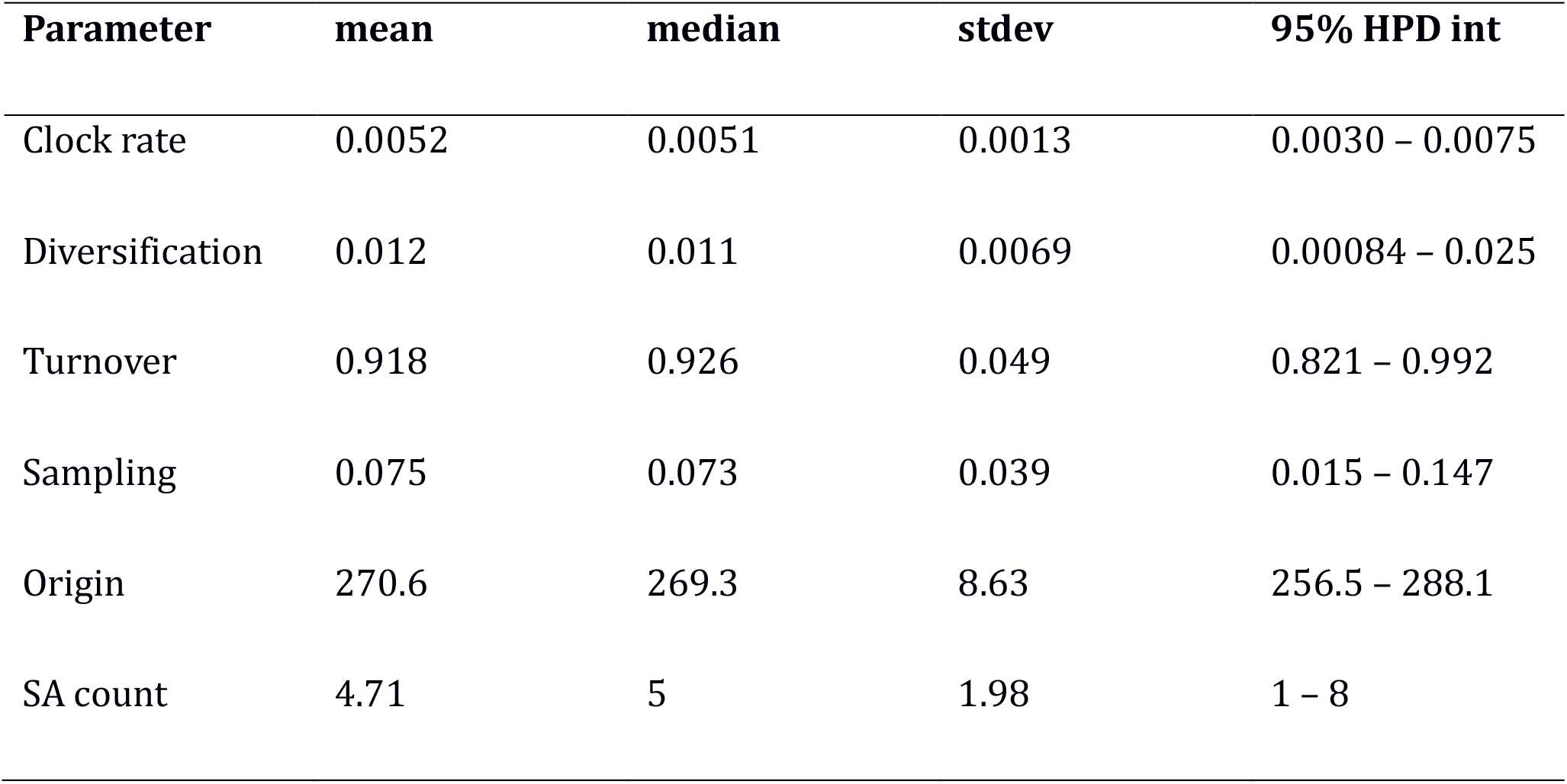
Results of the parameter estimates. Abbreviations: stdev = standard deviation, HPD int = highest posterior density interval, SA = sampled ancestors.

| Parameter | mean | median | stdev | 95% HPD int |
| --- | --- | --- | --- | --- |
| Clock rate | 0.0052 | 0.0051 | 0.0013 | 0.0030 – 0.0075 |
| Diversification | 0.012 | 0.011 | 0.0069 | 0.00084 – 0.025 |
| Turnover | 0.918 | 0.926 | 0.049 | 0.821 – 0.992 |
| Sampling | 0.075 | 0.073 | 0.039 | 0.015 – 0.147 |
| Origin | 270.6 | 269.3 | 8.63 | 256.5 – 288.1 |
| SA count | 4.71 | 5 | 1.98 | 1 – 8 |

The topology of the maximum clade credibility tree (Fig. 2) reveals several well-supported clades, although there are uncertainties in several areas of the tree. The placement of *Palaeobelemnopsis sinensis* within the Aulacoceratida is moderately well supported (posterior probability = PP = 0.69) as sister to the *Atractites alpinus* and *Aulacoceras sulcatum* clade. Furthermore, there is high support for including the sinobelemnitids within the Belemnitida (PP = 0.79), although Sinobelemnitidae itself is paraphyletic. Within the Belemnitida, there is weak support for a group containing mostly Jurassic taxa traditionally recognized as part of the Belemnitina (PP = 0.53), including *Schwegleria feifeli, Passaloteuthis bisulcata, Acrocoelites oxyconus, Megateuthis gigantea*, as well as *Lissajousibelus harleyi*, which was so far of uncertain placement inside Belemnitida. Sister to this clade is a larger group containing the remaining belemnites (PP= 0.67). Within this latter clade, several subclades displayed relatively high support: the sister group relationship between *Holcobelus munieri* and *Calabribelus pallinii* (PP = 0.79), a large clade containing belemnitids that share a pseudoalvelous (PP = 0.98); the same clade with *Belemnopsis apiciconus* as sister group to the latter is moderately supported (PP = 0.64). The clade containing *Praeactinocamax plenus, Gonioteuthis quadrata, Belemnitella propinqua* and *B. mucronata* is only weakly supported (PP = 0.44). Other clades received low to moderate support. Among these, we recovered a monophyletic clade containing *Cylindroteuthis puzosiana, Aulacoteuthis ernsti*, and *Oxyteuthis brunsvicensis* (PP = 0.68), which formed a weakly supported monophyletic clade (PP = 0.25) together with *Duvalia* and *Dicoelites* (PP = 0.57). Furthermore, we recovered a polyphyletic *Neohibolites* with *N. minimus* forming a monophyletic clade with *Dimitobelus diptychus* (PP = 0.39) and *N. ewaldi* as sister to *Mesohibolites minaret* (PP = 0.2).

**FIGURE 2.**
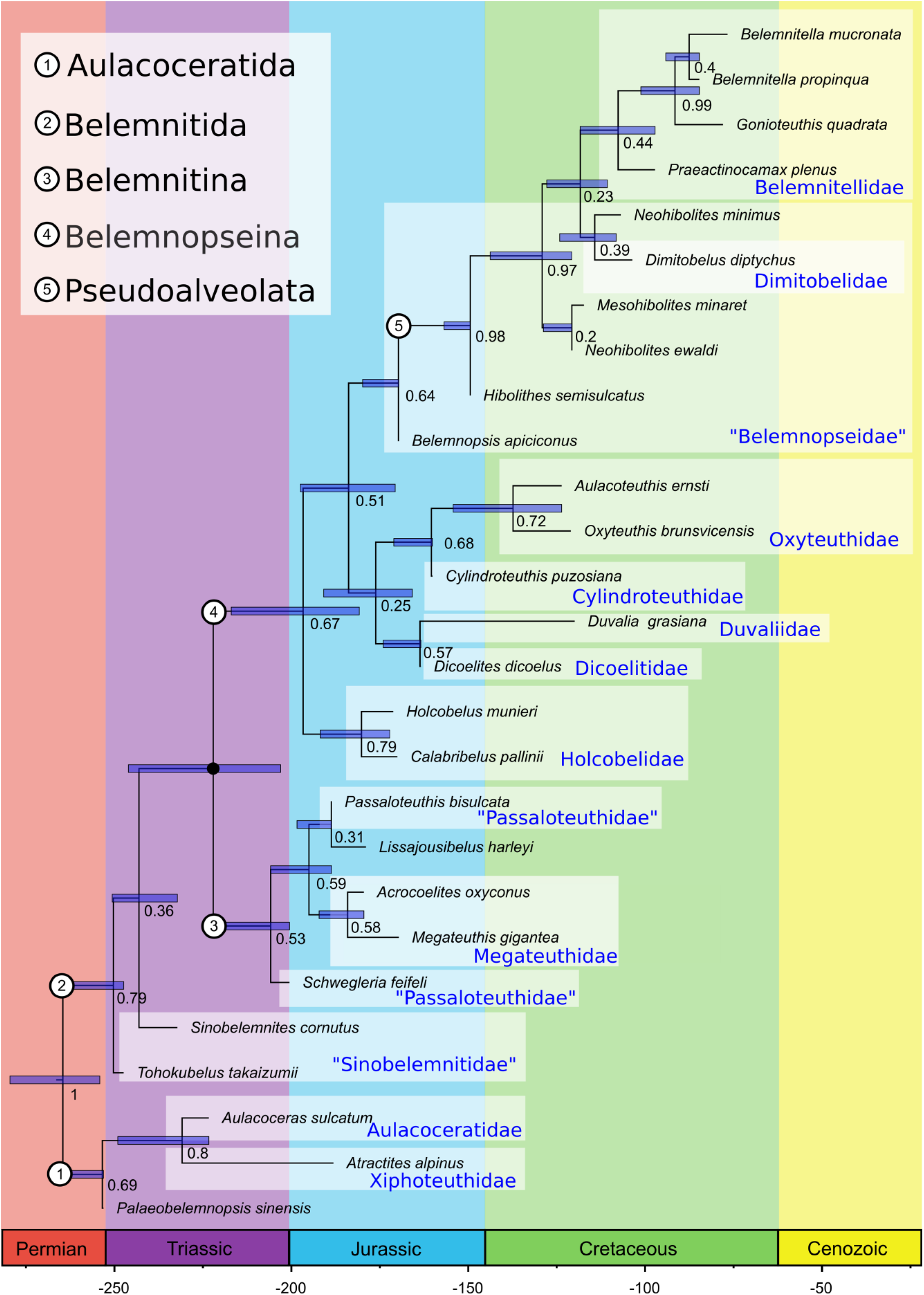
Maximum clade credibility tree of the Bayesian tip-dated analysis. Numbers at nodes represent posterior probability, while the blue bars indicate the 95% highest posterior density interval of the divergence time estimates. The small black dot represents the constrained clade. Tips with zero-length branches represent sampled ancestors.

## DISCUSSION

Due to the shared regeneration pattern and growth mode of Xiphoteuthidae (in the phylogeny represented by *Atractites alpinus*) and belemnites, Keupp and Fuchs (2014) suggested aulacoceratid paraphyly. On the other hand, Jeletzky (1966) argued for aulacoceratid monophyly, going so far as to view the group as an independent offshoot of bactritid cephalopods, leaving no descendants. Doyle et al. (1994) favored the derivation of belemnites from within the Aulacoceratida potentially via the Phragmoteuthida. Aulacoceratida are here recovered as a monophyletic group and sister to belemnites, with the Permian *Palaeobelemnopsis sinensis* confirmed as a member of the Aulacoceratida. However, since we only included a limited number of aulacoceratids in our analysis, it is still possible that total group Aulacoceratida is paraphyletic, also with respect to other groups of early coleoids such as the Phragmoteuthida.

Since Jeletzky (1966), all belemnites were usually divided into two suborders; Belemnitina and Belemnopseina, with members of the former group considered ancestral to the latter. Under this traditional scheme, Belemnitina groups taxa with apical furrows and Belemnopseina taxa with alveolar furrows. Problematic under the Belemnitina/Belemnopseina-scheme is that the earliest belemnites of the Sinobelemnitidae are considered to be Belemnopseina with a gap of ca. 25 Ma between the youngest Sinobelemnitida and remaining Belemnopseina (Iba et al., 2012). At least one of the two sinobelemnitids in the current analysis, on the other hand, forms the sister group to the remaining belemnites (Fig. 2). This result is in better agreement with the fossil record than the earlier hypothesis. It casts doubt on the homologization of dorsal alveolar furrows in belemnites, which was discussed as a potential uniting character of the Sinobelemnitidae with the Duvaliidae or Dicoelitidae (Iba et al., 2012). Still, a well-resolved position of the Sinobelemnitidae and clarification of their mono-or paraphyly will require a detailed study of this still sparsely known group and a better sampled Triassic fossil record of belemnites in general.

The species *Lissajousibelus harleyi* displays a ventral furrow in addition to dorsolateral apical furrows and was considered close to Belemnopseina by Weis et al. (2015b). In the present analysis, *L. harleyi* was found to be closely related to typical Belemnitina. Our Bayesian approach demonstrates how the relationship of belemnite taxa that do not easily fit the Belemnitina/Belemnopseina-scheme of Jeletzky (1966) can be resolved quantitatively by taking into account the maximally inclusive morphological and stratigraphic evidence. Belemnitina as defined herein is recovered as a monophyletic group with a likely origin in the Late Triassic but otherwise restricted to the Jurassic (Figs. 2 and 3). A close relationship between these taxa was already suggested by Stolley (1919), who grouped them into his family “Polyteuthidae”. This new definition includes at least the likely paraphyletic “Passaloteuthidae” and the Megateuthidae (Acrocoelitidae); it remains to be investigated in which way other Jurassic families not considered in the present analysis (e.g., Hastitidae, Salpingoteuthidae) are related to this group.

**FIGURE 3.**
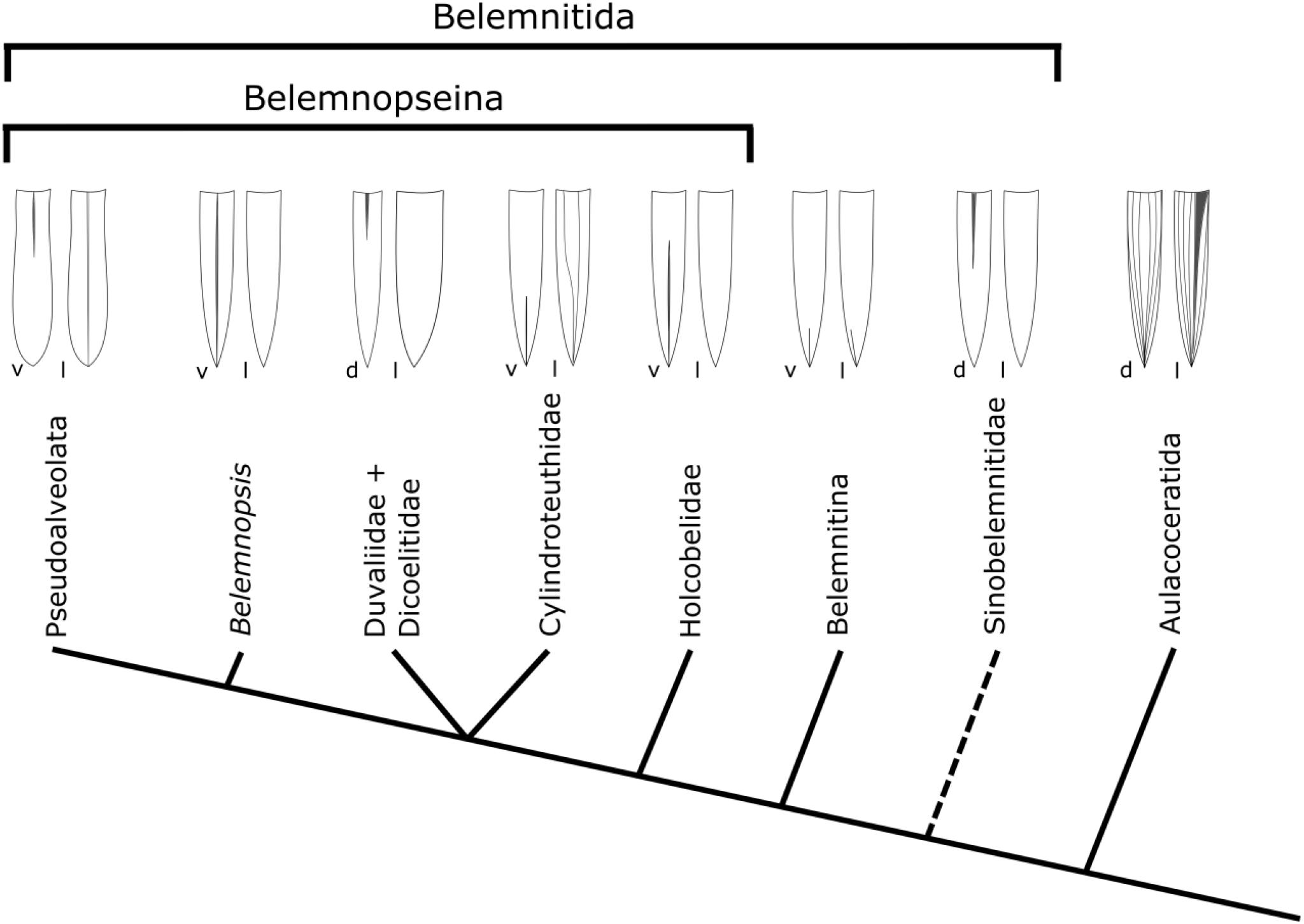
Cladogram showing the here suggested systematics of the Belemnitida based on the Bayesian tip-dated analysis. Sketches show the general outer morphological features of a typical representative of the groups in either dorsal (d), ventral (v), or lateral (l) view.

The Holcobelidae, Duvaliidae, and Dicoelitidae have an uncertain phylogenetic placement (e.g., Stolley, 1911a, 1911b; Stevens, 1964; Jeletzky, 1966; Combe morel, 1973; Weis et al., 2012). Our results place the Holcobelidae as the earliest branching Belemnopseina. This leaves it ambiguous whether their ventral furrow that does not reach the alveolus represents the ancestral state of the Belemnopseina or if this character represents a secondary development from an alveolar furrow reaching the alveolus as is seen in typical “Belemnopseidae”. In our tree, the placement of Duvaliidae as sister to Dicoelitidae finds moderate support, but the placement of this clade itself has only weak support (Fig. 2). The Duvaliidae share with the Dicoelitidae the presence of a dorsal alveolar furrow. However, a belemnopsein ventral alveolar furrow is also developed in the Dicoelitidae. *Duvalia grasiana* has recently been shown to display more organic-rich primary rostrum calcite than other belemnites (Stevens et al., 2022). A similar detailed description of the microstructure of other duvaliids and potentially related forms might reveal the phylogenetic position of this enigmatic group in future studies with more certainty.

The monophyletic clade containing the Oxyteuthidae as sister to *Cylindroteuthis puzosiana* confirms earlier thoughts on their phylogenetic relationships (e.g., Mutterlose, 1983). However, in contrast to these earlier hypotheses, we recovered the Cylindroteuthidae *sensu lato* clade within the Belemnopseina instead of within the Belemnitina. Still, the problem of the phylogenetic placement and evolution of Cylindroteuthidae and Oxyteuthidae needs further focused analyses, especially for Boreal belemnites of the Jurassic-Cretaceous transition, which will also have to include closely related species of the Pachyteuthidae. Our analysis suggests homology of the ventral furrow present in many Cylindroteuthidae *sensu lato*, not with the apical ventral furrow as is typical for the Megateuthidae/Acrocoelitidae, but with the belemnopsein ventral furrow. This hypothesis contrasts with the “stratophenetic” reasoning of the independent evolution of a ventral furrow in the genus *Aulacoteuthis* (Mutterlose and Baraboshkin, 2003; Baraboshkin and Mutterlose, 2004).

A name is suggested here for a newly identified and well-supported clade inside the Belemnopseina, the unranked Pseudoalveolata (see Appendix; PP=0.98; Fig. 2). The Pseudoalveolata is characterized by the synapomorphy of pseudoalveolus formation, a secondary alveolus-deepening, developing by dissolution/erosion of anterior organic-rich rostrum sections (Stevens et al., 2022). The pseudoalveolus is further characterized by a well-mineralized ‘“spike” projecting anteriorly toward the protoconch (*Nadelspitze* of Stolley, 1911a). *Hibolithes* has here been recovered as the earliest branching pseudoalveolate belemnite. Pseudoalveolus types have been considered of great importance in the phylogeny of the Belemnitellidae (e.g., Košťák, 2012), but contrary to suggestions by Dauphin et al. (2007) and Kosšťák and Wiese (2008), there is no conclusive evidence for the anterior rostrum of pseudoalveolate belemnites being of primarily aragonitic composition, as it likely consisted of calcite with primarily high organic contents (see Stevens et al., 2017; 2022).

The two species of *Neohibolites* analyzed herein (*N. ewaldi* and *N. minimus*) are not recovered as sister species. This indicates the possibly polyphyletic or paraphyletic nature of *Neohibolites*, a genus which has long been seen as ancestral to the two only belemnite families left after the Cenomanian, the Belemnitellidae and Dimitobelidae, (e.g., Mutterlose, 1998). Future studies focusing on the origin of Belemnitellidae and Dimitobelidae might shed more light on the exact origins of these two last surviving belemnite groups and their origins in the paraphyletic “Belemnopseidae”. “Belemnopseidae” has long been recognized as paraphyletic with respect to the Belemnitellidae and Dimitobelidae (e.g., Jeletzky, 1966; Mutterlose, 1988). However, a new phylogenetic definition of the family containing all belemnites closer to *Belemnopsis* than to the Pseudoalveolata seems possible. This would require a more thorough sampling of the diverse “belemnopseid” taxa of the earliest Cretaceous and a thorough revision of not only the genus *Belemnopsis* but also *Hibolithes*, encompassing detailed revision of the diverse *Belemnopsis* species of the Austral Realm (e.g., Stevens, 1965; Challinor, 1990). Based on the unclear and absent types for both genera (e.g., Combe morel and Howlett, 1993; Mitchell, 2015), the assignment of species to either genus was often based on superficial morphological assessment. We here confirm that there is no pseudoalveolus formation in the type species of *Belemnopsis, B. apiciconus*. Pseudoalveolus formation had already been suggested as a differentiating character of the genera *Belemnopsis* and *Hibolithes* by Stolley (1911a) but was unfortunately not followed on by later authors.

The tip-dated Bayesian analysis confirms earlier ideas that the epirostrum, a “tertiary” rostrum formation (Fuchs, 2012), which is developed only in some belemnites, represents a parallelism and does not indicate a close relationship (Bandel and Spaeth, 1988; Stevens et al., 2017). Epirostra are present in the analyzed species *Megateuthis gigantea, Holcobelus munieri, Calabribelus pallinii*, and *N. minimus*, found on disparate parts of our tree (Fig. 2).

In our proposed systematic framework (Fig. 3), we regard the potentially paraphyletic Sinobelemnitidae as the earliest branching belemnites. The remainder of the belemnites still falls into two large monophyletic clades, the Belemnitina and Belemnopseina, to preserve the current taxonomy as far as possible. We consider the Cylindroteuthidae *sensu lato*, Duvaliidae, and Dicoelitidae more derived than the Holcobelidae inside the Belemnopseina but otherwise of uncertain position with regard to the *Belemnopsis* + Pseudoalveolata clade.

The presented topology represents only a first step towards a well-resolved phylogeny of all belemnites. To achieve further resolution, it will likely be necessary to detect and evaluate further microstructural and geochemical data of several belemnite taxa and incorporate more taxa and characters into the analysis. Furthermore, well-preserved specimens from Konservat-Lagersta tten may provide additional valuable insights into the variability of soft-part anatomy, statoliths, radulae, hooks, or jaws within belemnites (e.g., Klug et al., 2010; Fuchs and Hoffmann, 2017), potentially adding other characters to include into phylogenetic analysis. This would also allow for a more inclusive phylogenetic analysis involving non-rostrum bearing “belemnoids” to resolve the decabrachian crown and stem groups.

## CONCLUSIONS

The first tip-dated analysis of belemnite (Belemnitida) phylogeny is presented. Our results suggest that the usually applied dichotomous subdivision of all belemnites into Belemnitina and Belemnopseina based only on the presence of apical *versus* alveolar furrows needs some adjustment. We consequently suggest the subdivision of all belemnites, except the potentially paraphyletic and early branching Sinobelemnitidae, into newly phylogenetically defined Belemnitina and Belemnopseina. Holcobelidae are the earliest branching Belemnopseina, Duvaliidae and Dicoelitidae are confirmed as Belemnopseina but are still of uncertain placement inside this group. A major change involves the transfer of the Cylindroteuthidae (including Oxyteuthidae) from the Belemnitina to the Belemnopseina. A new well-supported subgroup of the Belemnopseina, the unranked Pseudoalveolata, is suggested here, including a phylogenetic definition and the suggestion of a potential synapomorphy.

Because of their high fossilization potential and often high abundance in the marine fossil record, belemnites are particularly important in tracking faunal changes of the Jurassic and Cretaceous pelagic realms. The present study is only a first step; further analyses based on more taxa and characters, including detailed microstructural analyses of the rostra, are needed to further resolve more details of the belemnites’ evolutionary history. Applying quantitative and reproducible phylogenetic methodology in contrast to earlier approaches relying on authoritative overemphasis of single characters will lay a solid foundation for the future study of belemnites.

## APPENDIX Phylogenetic Definition of Pseudoalveolata

**Pseudoalveolata unranked K. Stevens, A. Pohle, and R. Hoffmann, new clade name.**

Phylogenetic definition: the least inclusive clade containing the Belemnitellidae Pavlow, 1914, Dimitobelidae Whitehouse, 1924, and the species *Hibolithes semisulcatus* (zu Mu nster, 1830), *Neohibolites ewaldi* (Strombeck, 1861), *N. minimus* (Miller, 1826), and *Mesohibolites minaret* (Raspail, 1829). Etymology: derived from the belemnite morphological term pseudoalveolus, itself derived from Greek pseu do, meaning “to lie” or “to deceive”, and alveolus, from Latin, meaning a small cavity. Reference phylogeny: Fig. 2. Diagnosis: In belemnite morphological terminology, the alveolus is the approximately cone-shaped cavity of the rostrum, which contains the phragmocone. If the alveolus is secondarily enlarged by abrasion or dissolution of the anterior part of the rostrum due to an anterior primarily porous and organic-rich composition of the rostrum (e.g., Stolley, 1911a; Ernst, 1964; Stevens et al., 2022), the resulting secondary deepening is termed a pseudoalveolus. Anterior primarily porous and organic-rich rostra, which might result in a pseudoalveolus, accordingly represent the synapomorphy of the Pseudoalveolata.

