## Supplementary material for "Bayesian inference reveals a complex evolutionary history of belemnites": Supp File 1

### Supplementary File 1

#### Character description

Character 1: Two dorsolateral apical furrows absent (0) or present (1).

Character 2: Apical ventral furrow absent (0) or present (1).

Character 3: Two ventrolateral apical furrows absent (0) or present (1).

Character 4: Ventral alveolar furrow(s) absent (0) or present (1).

Character 5: Ventral alveolar furrow(s) intermediate (0) or reaching alveolus (1).

Character 6: Ventral alveolar furrow number 1 (0) or 2 (1).

Character 7: Ventral furrow reaches from alveolus towards apical region no (0) or yes (1).

Character 8: Ventral splitting surface (slit, slitfield, “Schlitzfeld”) absent (0) or present (1).

Character 9: Ventral splitting surface type rudimentary (0) or full (1)

Character 10: Dorsal alveolar furrow absent (0) or present (1).

Character 11:  Dorsal slit (splitting surface, slitfield, “Schlitzfeld”) absent (0) or present (1).

Character 12: Doppellinien absent (0) or present (1).

Character 13: Doppellinien parallel (0), diverge towards alveolus (1), or multiple (2).

Character 14: Rostrum proper aragonite (0) or calcite (1).

Character 15: Pseudoalveolus, secondary erosion of alveolus forming a concave or conical structure, absent (0) or present (1).

Character 16: Pseudoalveolus surface convex (0) or concave (1)

Character 17: Alveolus angle low <~11° (0) or high >~11° (1).

Character 18: Alveolus central (0) or ventrally displaced (1).

Character 19: Rostrum ribbed with longitudinal ridges (0) or smooth (1).

Character 20: Two dorsolateral longitudinal depressions absent (0) or present (1).

Character 21: “Vascular” imprints absent (0) or present (1).

Character 22: Apex type acute (0) or obtuse (1).

Character 23: Epirostrum absent (0) or present (1).

Character 24: Cameral deposits absent (0) or present (1).

Character 25: Type of juvenile rostrum conirostrid (0) or clavirostrid (1).

Character 26: Rostrum hastate in dorsal view no (0) or yes (1).

Character 27: Deep lateral grooves absent (0) or present (1).

Character 28: Cross-section at position of protoconch +/- circular (0) or compressed (1).

Character 29: Ventral tongue-like extension absent (0) or present (1).
